# MGX 2.0: Shotgun- and assembly-based metagenome and metatranscriptome analysis from a single source

**DOI:** 10.1101/2023.09.21.558800

**Authors:** Sebastian Jaenicke, Sonja Diedrich, Alexander Goesmann

## Abstract

Metagenomics studies have enabled scientists to analyze the genetic information of natural habitats or even complete ecosystems, including otherwise unculturable microbes. The processing of such datasets, however, remains a challenging task requiring extensive computational resources. MGX 2.0 is a versatile solution for the analysis and interpretation of microbial community sequence data. MGX 2.0 supports the processing of raw metagenomes and metatranscriptomes, but also enables assembly-based strategies, including downstream taxonomic binning, bin quality assessment, abundance quantification, and subsequent annotation coming from a single source. Due to the modular design of MGX, users are able to choose from a wide range of different methods for microbial community sequence data analysis, allowing them to directly compare between read-based and assembly-based approaches or to evaluate different strategies to analyze their data.

## Introduction

Metagenomics, the sequence-based study of microbial communities, has emerged as a popular scientific field. Facilitating access to the genetic information of unculturable microbes, the technique is widely applied to study the composition of microbial consortia and the metabolic potential encoded in their genomes. Metagenomics studies have enabled scientists to gain a deeper understanding of the functioning of natural environments^1, 2^, identify key factors that enable synthetic microbial assemblages to perform a common task^3, 4^, or determine correlations between microbiome composition in human-associated habitats and the prevalence of certain diseases^5^ or human health in general^6^. While metabarcoding approaches provide high resolution based on a single marker gene such as that encoding for 16S rRNA at comparably low cost, they only allow taxonomic characterization of a community to be performed.

Whole metagenome shotgun sequencing, on the other hand, gives researchers access not only to taxonomic profiles of a microbial population, but also allows deduction of the metabolic capabilities of its members based on the characterization of genes and gene fragments. Nowadays, researchers are able to choose from a large variety of specialized tools for the analysis of environmental sequence data. Software packages such as Kraken 2^7^, Kaiju^8^, or Centrifuge^9^ enable the high-throughput taxonomic classification of short microbial sequences. For the functional annotation of genes and gene fragments, alignment-based approaches prevail; for this purpose, database comparisons are performed with advanced homology search algorithms such as DIAMOND^10^ or HMMER^11^ using either six-frame-translations of the nucleotide query sequences as an input, or presumably protein-coding sequences after a preceding gene prediction step^12, 13^.

With the ever-increasing output of current sequencing instruments, there is also a trend towards the generation of larger metagenome datasets that offer more complete coverage of the studied microbial communities. These metagenomes comprise sufficient sequence information to perform metagenome assembly, enabling the reconstruction of full-length genes, identification of gene clusters, and ultimately, the recovery of metagenome-assembled genomes (MAGs), which represent consensus genomes of closely related microbial strains. In previous studies^14, 15^, metagenome assembly has successfully been applied to recover thousands of genomes from different microbiomes. Provided they are sufficiently large, datasets may either be assembled individually, or several metagenomes from closely related environments are co-assembled jointly in order to increase overall coverage and to improve subsequent taxonomic binning. For this purpose, specialized *de novo* metagenome assemblers such as MEGAHIT^16^ or metaSPAdes^17^ have been developed, that specifically address the challenges inherent to metagenome sequence data such as non-uniform read coverage. After the assembly step, contigs are assigned to different taxonomic bins based on intrinsic properties such as read coverage or nucleotide composition^18, 19^. The quality of the resulting bins is then assessed with *e.g.* CheckM^20^ or BUSCO^21^, determining the levels of completeness and possible contamination based on the occurrence of lineage-specific marker genes.

Finally, transcriptome sequencing (RNA-Seq) is commonly applied to identify the regions of a genome that are actively transcribed. Traditionally executed by reverse-transcribing RNA into cDNA, which is then sequenced, recent approaches such as nanopore sequencing also allow direct sequencing of the individual RNA molecules^22^, thereby avoiding potential biases inflicted by the reverse transcription step. While gene expression cannot be directly quantified with this approach, transcriptional changes may be derived by contrasting datasets obtained under different conditions. To this end, sequence data is aligned to the annotated genome and reads are allocated to corresponding genes based on the alignments’ location^23^. In the absence of an appropriate reference genome, the transcriptome of an organism may also be reconstructed employing *de novo* transcriptome assemblers such as Trinity^24^, or rnaSPAdes^25^.

Applied to a microbial population, transcriptome sequencing (metatranscriptomics) enables distinction between active community members and those that are dormant, identification of possible symbiotic relationships between organisms, and afford valuable insights into community alterations caused by external stimuli such as *e.g.* environmental changes^26^. Given that corresponding reference genomes are typically not available, and in order to obtain an unbiased representation of the active microbial community, metatranscriptome data can either be subjected to read-based analysis, or *de novo* assembly might be attempted. No significant progress has recently been made towards the development of dedicated metatranscriptome assemblers, and the sole published software package for this purpose, IDBA-MT^27^, has been demonstrated to achieve suboptimal results^28^. However, several studies have favorably evaluated the application of *de novo* transcriptome assemblers to metatranscriptome datasets^26, 28^.

Regardless of whether a read-based or assembly-based approach is chosen, significant compute resources are required for the timely analysis of large-scale microbial community datasets. In addition, software for this purpose is almost exclusively provided in the form of command-line-based tools for the Linux operating system. The processing of unassembled metagenomes and metatranscriptomes, to a certain degree, is easily parallelized and can therefore be distributed to compute infrastructures such as HPC clusters or cloud-based resources. Resource requirements for assembly are drastically larger, necessitating access to compute nodes with large CPU core numbers as well as generous amounts of system memory. Subsequent annotation of predicted genes, however, is eased for assembled data due to the reduced number of input sequences in comparison to unassembled datasets, and relative abundances can be obtained by aligning the raw input data to the assembled contigs.

Established metagenomics platforms such as MG-RAST^29^, IMG/M^30^, or EBI MGnify^31^ provide large-scale compute resources for the analysis of environmental sequence data. MG-RAST focuses on unassembled metagenome datasets, but also supports the submission of preassembled contigs. With IMG/M, read-based metagenome analysis is no longer offered as a public service; since 2017, submissions from outside the JGI have been restricted to preassembled data^32^, requiring users to procure appropriate compute resources for the initial assembly step. EBI MGnify^31^ offers to perform assembly of individual metagenomes, but necessitates sequence data to be publicly deposited in the European Nucleotide Archive (ENA) beforehand. Command-line workflows such as CoMW^33^ or SAMSA2^34^ specifically address metatranscriptome analysis, but have been implemented for either local execution or “for standalone use on a supercomputing cluster”^34^. Given the widespread availability of high-throughput sequencing and the constantly rising number of metagenomics studies being conducted, there is a clear demand for modern metagenomics analysis platforms offering access to state-of-the-art tools and compute resources; this is especially important for researchers without computational expertise or without access to appropriate compute infrastructure.

With the initial release of the MGX framework^35^, we previously introduced a highly flexible solution for the management and analysis of unassembled metagenome datasets. MGX features a large selection of analysis workflows addressing taxonomic classification as well as functional profiling, antibiotic resistance gene screening, and reference genome alignment. As a client/server solution, no extensive compute resources need to be provisioned by its users. MGX supports confidential projects for the analysis of unpublished data; however, we encourage the submission of raw sequence data to the appropriate INSDC repositories upon publication in order to facilitate the reproducibility of results as well as enable potential re-analysis. Since its publication, MGX users have successfully applied the software to process metagenomic datasets originating from a large variety of environments, including soil^2, 36, 37^, aquatic^38^, and synthetic^4^ habitats, allowing researchers to gain valuable insights into the respective microbial populations found in these ecological niches.

Up until now, no software application has addressed both metagenome co-assembly and *de novo* metatranscriptome assembly, and in the past users pursuing assembly-based analyses had to perform these manually and without the possibility to conveniently compare results to those obtained from read-based analyses. Here, we describe the most recent extension of the MGX framework, MGX 2.0, which now also incorporates support for the assembly of microbial community sequence data followed by subsequent taxonomic and functional annotation.

## Results

### MGX 2.0 enables metagenome and metatranscriptome assembly

With its most recent extensions, the MGX 2.0 platform introduces novel and reproducible workflows for the assembly of whole-metagenome shotgun datasets as well as metatranscriptome assembly and a large variety of different pipelines targeting the analysis of predicted genes. To achieve this, we have significantly extended the application and its data model, thus enhancing its capabilities. Among others, out novel workflows include steps for quality control, assembly, and gene prediction. Taxonomic binning is performed for metagenomic assemblies, while an additional step of the metatranscriptome processing pipeline performs the removal of ribosomal sequences prior to assembly. By realigning raw reads to the assembled contigs and transcripts, read coverage is obtained, which can be exploited to directly compare read and assembly-based analysis results.

### Analysis workflows for taxonomic and functional annotation of microbial community sequencing data

For MGX, we previously developed a large variety of workflows employing tools for the taxonomic and functional characterization of metagenome datasets, that we continuously extended using novel algorithms and databases. We now have complemented these with comprehensive analysis pipelines targeting the annotation of assembled metagenome and metatranscriptome data. For this purpose, we implemented a large variety of novel workflows to annotate predicted genes with, among others, COG groups, EC numbers, KEGG KOfams, or Pfam protein domains (Suppl. Tables 1-3). These workflows are provided with predefined parameters, but users are free to adapt *e.g.* cutoff values based on their needs.

### Extensions of the MGX 2.0 user interface

Novel components of the MGX 2.0 graphical user interface enable scientists to comfortably execute analyses, visualize the corresponding results upon job completion, and export data in a variety of formats. The “blob plot” component provides an initial overview of an assembly, displaying contigs or transcripts based on size, GC content, coverage, and taxonomic assignment (Fig. 1). Assembled contigs can be evaluated in detail using the Bin Explorer, which interactively displays taxonomic and functional classifications assigned to the predicted genes (Fig. 2) and allows users to search for annotations of interest. Most notably, assembly support has been seamlessly integrated into the MGX 2.0 attribute visualization component, thereby allowing direct comparisons to be made between unassembled and assembled data (Fig. 3) taking read coverage into account.

**Figure 1:**
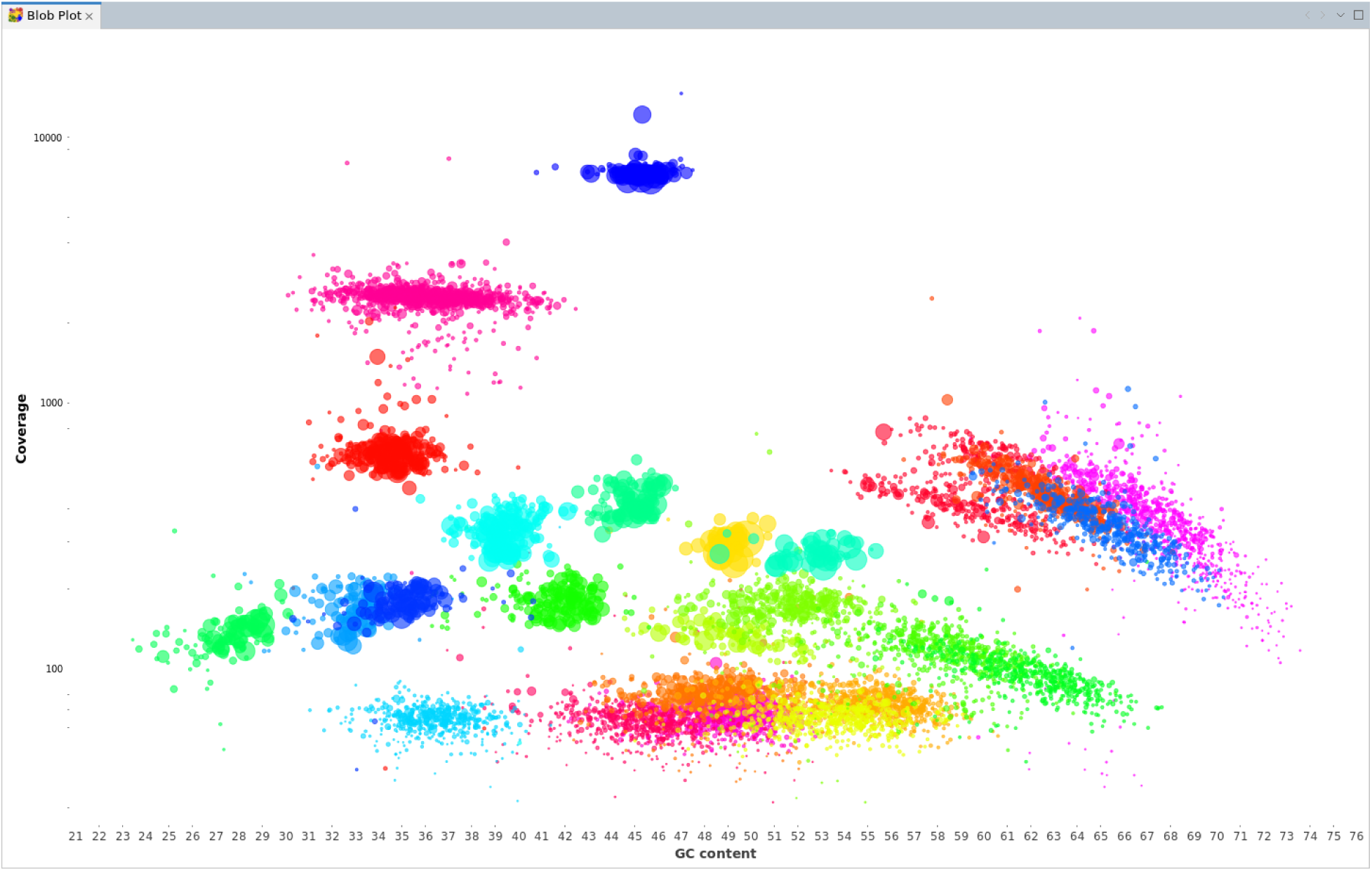
Metagenome assembly overview. The MGX “blob plot” provides an overview of one or several assembled metagenomic bins, allowing to visually identify contigs with similar properties. Based on GC content and read coverage, metagenomic contigs are displayed as circles of different sizes depending on the contigs’ length, and colored according to the taxonomic assignment of the metagenome bin.

**Figure 2:**
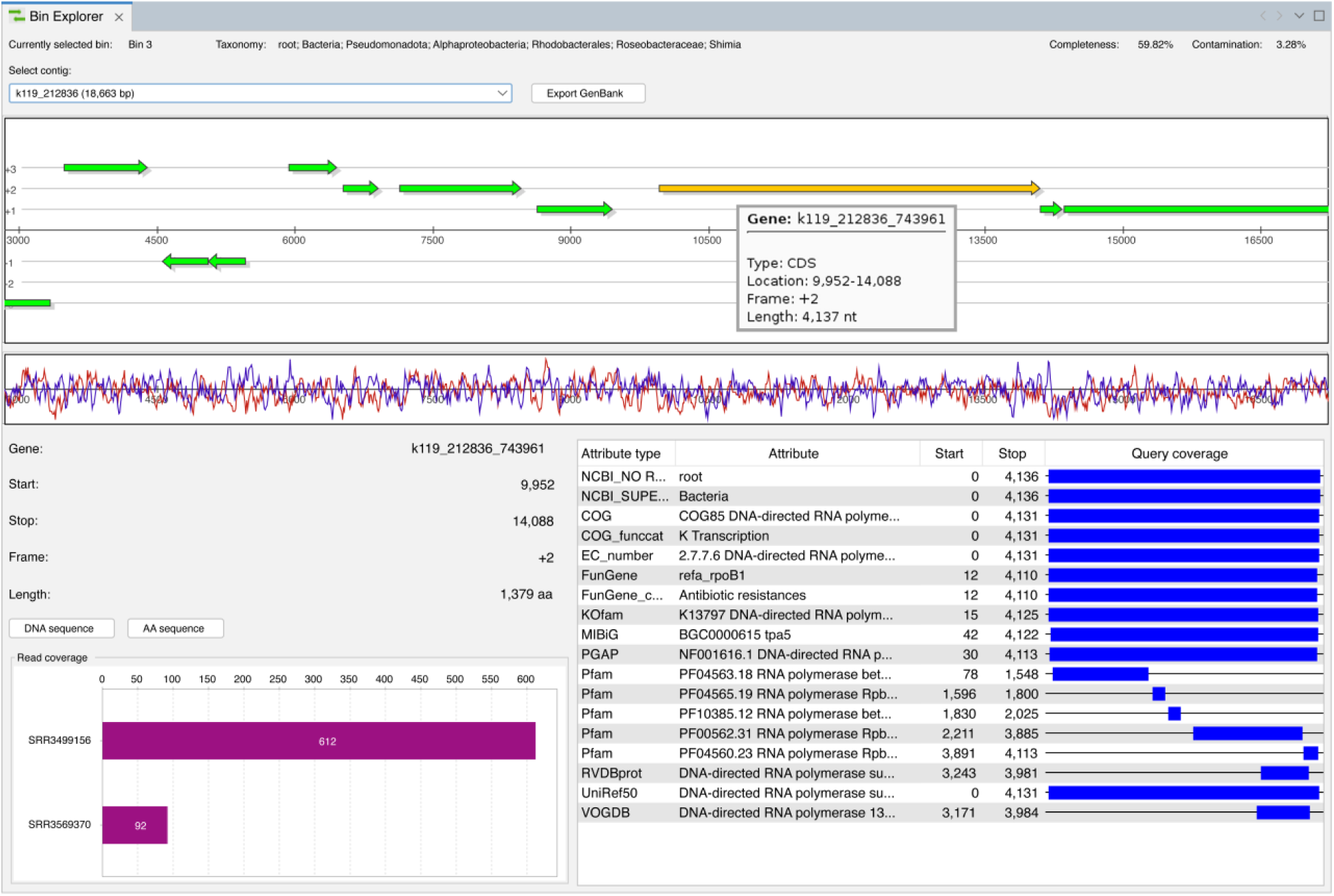
Interactive assembly inspection. The “Bin Explorer” facilitates interactive exploration of assembled contigs and associated annotations. For each contig, a graphical depiction of the contig and its predicted genes is provided. Once a gene is selected, additional details about the location of the gene, its predicted taxonomic origin, and functional features that were detected are displayed. An additional coverage chart (bottom left) provides abundance information, *i.e.* the number of metagenome reads in co-assembled datasets that overlap with the selected gene.

**Figure 3:**
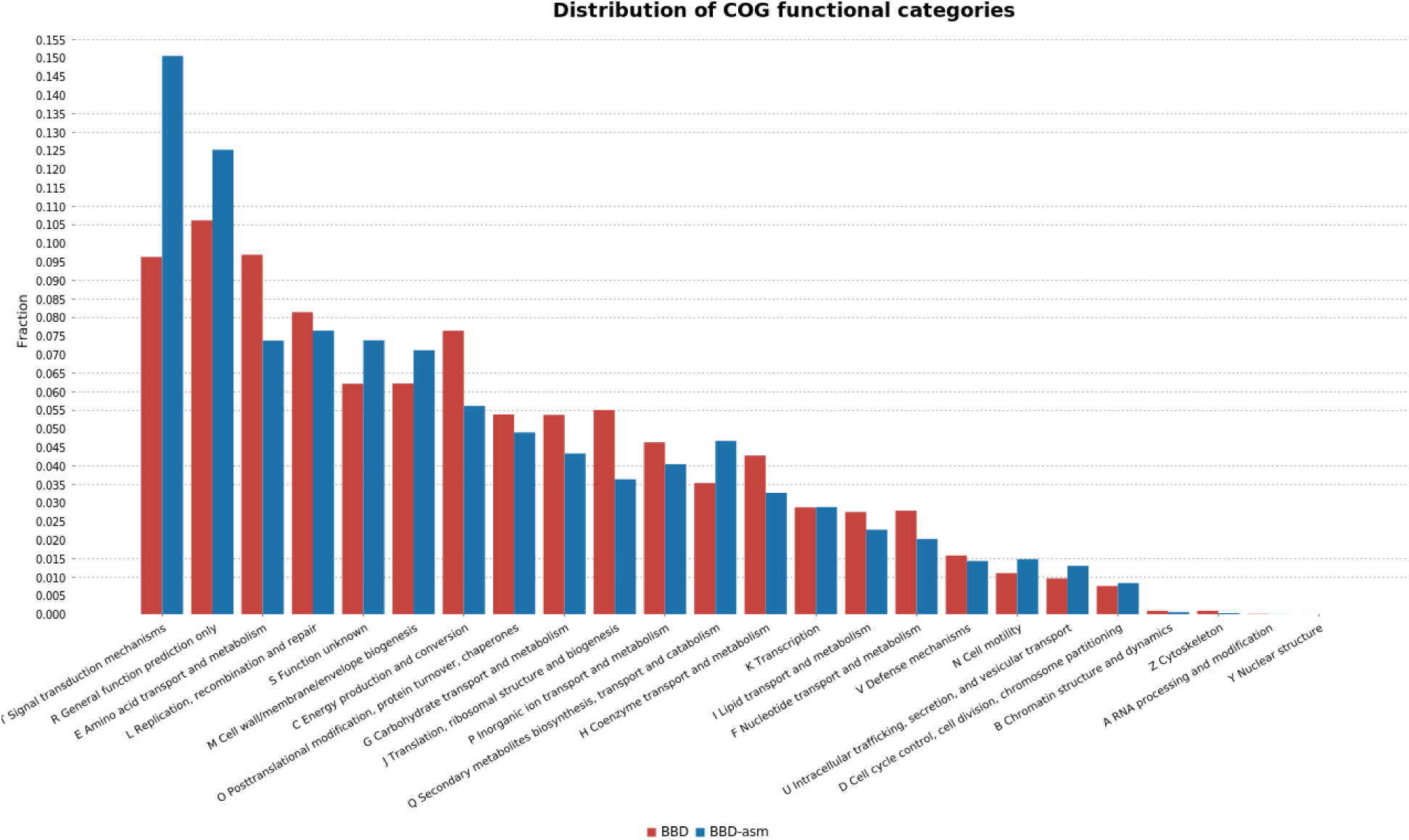
Joint functional annotation of microbial community data. Assembly support has been seamlessly integrated into MGX 2.0, enabling users to perform direct comparisons between the results of read and assembly-based analyses. For the BBD metagenome, *e.g.* annotated functional COG categories are easily compared between read-based (BBD; red) and assembly-based (BBD-asm; blue) strategies.

### Improved re-analysis of public metagenome and metatranscriptome datasets

Based on the re-analysis of a dataset targeting black band disease (BBD) in corals^39^, we demonstrate the advantages of the MGX 2.0 platform. Applying a co-assembly strategy of BBD and cyanobacterial patch (CP) metagenomes, MGX computed a more complete and representative assembly; while Sato *et al.* reported upon 19 bins (5 of them >90% complete), MGX recovered 25 bins (15 bins with >90% completeness; Suppl. Table 4), among them one genome classified to be complete (*Ruegeria arenilitoris*; 100% completeness, 0.43% contamination). In the original study, 48.3 Mbp comprising 50,617 genes were assigned to bins; the assembly workflow within MGX 2.0 was capable of binning significantly more contigs (8,883 contigs; 89.2 Mbp) and genes (86,092).

Of special interest are two bins of cyanobacteriotal origin, that represent metagenome-assembled genomes (MAGs) of dominant microbial community members: Bin 2, which was almost exclusively assembled from the CP metagenome and assigned to *Okeania sp. KiyG1*, and bin 24, which mainly originates from the BBD metagenome and was classified as *Roseofilum reptotaenium*. We assume these to be equivalent MAGs to the bins Cya1 and Cya2 reported by Sato *et al.* Especially for bin 2, completeness (91.16%; previously 60.2%) and contamination estimates (2.4%; previously 4.3%) improved significantly over the previous results. Since the authors allocated Cya1 to *Trichodesmium erythraeum*, we independently confirmed the MGX 2.0 assignment of bin 2 to *Okeania sp. KiyG1* with GTDB-Tk, which assigned it to the *Okeania* genus (Suppl. Table 5), and with the EDGAR phylogenomics application^40^(Suppl. Fig. 1). Also, read-based analyses within MGX did not confirm the presence of *Trichodesmium* in the CP metagenome; instead, *Okeania* is identified as the dominant genus (34.6%), followed by *Ruegeria* (10.4%).

*Roseofilum reptotaenium*, which is represented by bin 24, has previously been described as being causative for BBD lesions^41^. *Roseofilum* represents the most abundant genus (79.5%) in the unassembled BBD metagenome, followed by *Alteromonas* (2%). The MAG, comprised of 108 contigs with 4,869 predicted genes, corresponds to the Cya2 bin described by Sato *et al.*; according to the CheckM assessment within MGX, the recovered bin is 99.11% complete with 0.22% remaining contamination. Interestingly, MetaPhlAn 4, a commonly applied taxonomic profiling tool, only assigned small numbers of reads to the Cyanophyceae class for the BBD metagenome (3,447 reads; 6.57%), and completely failed to identify *Roseofilum* at the genus level in both metagenome and metatranscriptome data (Suppl. Fig 2). Applying the MGX default classification pipeline or the Metabuli workflow within MGX 2.0 to the BBD metagenome, *Roseofilum* was identified as the most abundant genus by both algorithms.

In total, the novel assembly workflow in MGX 2.0 predicted 196,800 genes for the complete metagenome co-assembly, with 110,708 genes not being allocated to any metagenomic bin. The metatranscriptome assembly yielded 180,779 transcripts with 192,859 predicted genes. As reported previously, functional analysis of the raw and assembled metatranscriptomes confirmed high transcriptional activity especially for genes involved in photosynthesis, *e.g.* photosystems I and II (EC: 1.97.1.12 and 1.10.3.9); phenylalanine ammonia-lyase (EC: 4.3.1.24) and ribulose-bisphosphate carboxylase (EC: 4.1.1.39) were found to be highly enriched especially in the BBD metatranscriptome. Applying the corresponding workflows within MGX 2.0, a more comprehensive and complete analysis of the available sequence data was thus achieved.

## Discussion

MGX (https://mgx-metagenomics.github.io/) is a client/server framework for the management and analysis of microbial community datasets such as metagenomes and metatranscriptomes. MGX features a large selection of highly specific analysis workflows providing access to novel tools for taxonomic classification and functional assignment of unassembled metagenomic datasets. With MGX, metagenomic sequences are easily processed and annotated with COG groups, EC numbers, KEGG pathways, protein domains, or antibiotic resistance genes. MGX also supports the alignment of metagenome and metatranscriptome sequences to user-provided reference genomes, thereby enabling the creation of fragment recruitment plots or visual identification of expressed genes and gene clusters.

With its most recent extension, the MGX 2.0 platform now provides support for the assembly of whole-metagenome shotgun datasets as well as metatranscriptome assembly, followed by subsequent annotation of assembled contigs or transcripts. For metagenomes, a sophisticated state-of the-art workflow is provided, addressing quality control, metagenome assembly, taxonomic binning, abundance calculation, and finally, gene prediction. Bin completeness and pureness assessment enable users to evaluate the success of the assembly and binning procedures implemented within MGX 2.0.

For metatranscriptome data, a similar approach is being followed, but with an additional filtering step in order to improve the quality of the assembly. No taxonomic binning step is carried out for metatranscriptome assemblies, since neither read coverage nor sequence composition of assembled transcripts can be used for this purpose. No algorithm is currently available that facilitates the taxonomic binning of metatranscriptomes, and the development of appropriate methods is a task open for future work.

Additional workflows offered within MGX 2.0 facilitate the taxonomic and functional annotation of predicted genes residing on the assembled contigs and transcripts. Similar to those already available for the analysis of unassembled datasets, these workflows are able to annotate predicted genes with different functional categorization schemes such as COGs, KOfams, or EC numbers, identify conserved domains or perform high-throughput screening for gene families of interest. All these workflows come with predefined parameters, but the interactive analysis wizard within MGX 2.0 enables researchers to easily adapt them to their own needs.

Since assembly represents a non-linear filtering step, it is normally not possible to directly compare results obtained from read-based analyses with annotations derived from assembled contigs. Therefore, MGX 2.0 keeps track of the number of reads overlapping with each predicted gene for each dataset and is hence able to reconstruct abundance information. Within the MGX 2.0 graphical user interface, novel components allow interactive inspection of the outcome of an assembly, visualization of annotations, comparisons of unassembled and assembled metagenomes and –transcriptomes, and data export *e.g.* for statistical assessment.

The advantages of MGX 2.0 are demonstrated via the re-analysis of public metagenome and metatranscriptome datasets from a study focusing on black band disease in corals. The MGX 2.0 assembly workflow was able to drastically improve over the results obtained previously, recovering more high-quality MAGs and assigning almost twice as many genes to taxonomic bins. Also, the option to execute different algorithms for taxonomic classification has to be considered an important advantage of MGX 2.0 over static workflows that rely on only one method; in this case, *Roseofilum*, the most abundant genus in the BBD datasets and likely the root cause for the coral lesions, was completely missed by a popular tool.

Functional analysis results obtained for the metatranscriptomes largely confirmed the results obtained in the previous study, identifying highly transcribed photosynthesis genes in both datasets. With MGX 2.0, scientists have access to a modern set of different tools for the analysis of microbial community sequence data that is unmatched by other platforms. Until now, common approaches like co-assembly or taxonomic binning were not offered within a metagenomics application and had to be performed manually. As its rich feature set supports both read-based as well as assembly-based strategies, convenient data visualizations, and various export options, users are enabled to pursue their study goals without being dependent on knowledge of the Linux command line or access to large-scale compute infrastructures.

## Methods

### Extension of the MGX 2.0 infrastructure

In order to support metagenome and metatranscriptome assembly, the MGX data model was significantly extended in comparison to the initial MGX release and several adaptations had to be made, *e.g.* to properly track single-end and paired-end datasets. Information about the assembly, bins, contigs, and predicted genes is retained within the PostgreSQL database associated with each MGX project (Suppl. Fig. 3). We also established a new REST-based interface for data retrieval and entity creation, thereby abolishing the previous requirement of a shared file system and direct database access for the newly created assembly workflows; instead, data is transparently retrieved, processed by the workflow and results are then submitted to the MGX project database via the REST service. Following this approach, the execution of assembly workflows can also be distributed to *e.g.* cloud-based compute infrastructures.

### MGX 2.0 enlarges the portfolio of methods for raw metagenome analysis

Based on feedback and requests from users of the MGX application and our own internal evaluations, newly developed tools and databases were either continuously added or updated to the most recent versions. At the same time, support for several tools was abolished from MGX where the method can no longer be considered to be state-of-art or where recent database releases are no longer offered. Currently, MGX 2.0 offers a broad and modern set of over fifty different pipelines for the taxonomic and functional analysis of raw and assembled metagenomes and –transcriptomes, incorporating tools and databases such as MetaPhlAn 4^42^, Metabuli^43^, MIBiG^44^, or PGAP^45^.

### CommonWL enables reproducible metagenome and metatranscriptome assembly

So far, MGX has exclusively relied upon the Conveyor workflow engine^46^ for sequence data processing. Conveyor streams data between the individual processing steps, only creating files when external programs are being invoked. While this approach has several performance advantages, it also means that large amounts of data are constantly kept in system memory, an approach rather impractical for the assembly of large datasets.

With the development of MGX 2.0, support for workflows implemented in the Common Workflow Language^47^ (CWL) has since been added. CWL was chosen as it represents an open workflow definition standard that is agnostic from a distinct implementation. Different CWL implementations are available that allow, in addition to local execution, the distribution of tasks to compute clusters (*e.g.* Grid Engine, Torque, or SLURM) or cloud infrastructures such as OpenStack, AWS, or Azure. By using modern containerization techniques, Docker images can be utilized to ensure the reproducibility of results. Within MGX, workflow parameters are automatically extracted from the corresponding definition file and offered to the user for possible adaptation.

### Metagenome assembly, binning, and abundance calculation

The newly established workflow for metagenome assembly has been designed to target the processing of Illumina sequence data as the currently predominant sequencing platform. While the inclusion of single-end sequencing data is supported in general, we require that at least one paired-end dataset is provided to improve assembly results. Also, only information from paired-end data is used during the taxonomic binning stage, and we recommend including multiple datasets, as this has previously been shown to improve binning efficiency^18, 48^. Sequences obtained with PacBio or Oxford Nanopore technology can be supplied as single-end datasets, but we advise performing prior read correction as these sequences are often prone to error^49, 50^. During workflow execution, sequence data is initially preprocessed using fastp^51^ in order to perform adapter removal as well as quality-based trimming. The metagenome assembly step itself is based on the MEGAHIT^16^ assembler in sensitive mode (Suppl. Table 6). For abundance estimation, metagenome sequences are mapped back to the assembled contigs employing Strobealign^52^ for alignment and SAMtools^53^ for data conversion. VAMB^54^, MetaBat 2^19^, and SemiBin 2^55^ are then applied for taxonomic binning; as VAMB bins every input sequence, its output is post-processed, filtering out bins less than 100 kbp in size. Binning predictions are combined into an optimized result using DAS Tool^56^, and bin completeness as well as contamination estimates are assessed via CheckM^20^. The obtained contigs are taxonomically classified using Metabuli, and taxonomic assignments of the bins are then derived from the classification results of the individual contigs contained therein, relying upon a majority-based scheme (https://github.com/sjaenick/assignBin/) that requires a fraction of at least 55% identical assignments on a taxonomic rank after prior pruning of low-abundant predictions. The complete assembly is then subjected to a gene prediction step employing a parallelized version of Prodigal^12^, and finally, abundance counts of genes and gene fragments in the datasets are determined via the featureCounts software^23^, while bamstats (https://github.com/sjaenick/bamstats/) is used to compute total read coverage of the contigs.

### Metatranscriptome preprocessing and assembly

The metatranscriptome assembly pipeline implemented within MGX 2.0 follows the same general approach as the metagenomics workflow. Initially, all datasets are preprocessed with RiboDetector to remove sequences of ribosomal origin, as these are often present in high abundance and due to their conserved subregions, are prone to contribute to misassemblies. The adapter autodetection of fastp^51^ is unsuitable for transcriptome data, since overrepresented sequences might be the result of high levels of transcription instead of actual sequencing adapters. Therefore, adapter clipping of single-end and paired-end sequences and quality-based trimming is performed with Trimmomatic^57^ instead. Metatranscriptome assembly is implemented via the rnaSPAdes software^25^, which was chosen due to its ability to recover more genes and gene fragments in comparison to Trinity^24^ in our internal evaluations. To predict genes on the assembled transcripts, Prodigal^12^ is used in its metagenomic mode. Afterwards, reads are realigned to the assembled transcripts with Strobealign^52^ and SAMtools^53^, and read coverage analysis of assembled transcripts and predicted genes is performed for each input dataset with featureCounts and bamstats, as described above.

Read coverage of assembled transcripts correlates with both the relative abundance of the source organism as well as transcriptional activity. In addition, the sequence composition of gene-coding regions differs from that of genomic contigs. Binning software from the metagenomics field thus cannot be employed, as these tools typically rely on both coverage information and sequence composition. So far, no method has been published to perform binning of assembled metatranscriptome transcripts; hence, no further processing is being performed and all assembled transcripts are assigned to one single bin, which denotes unbinned data.

### Novel standardized workflows for the annotation of MAGs and assembled metatranscriptomes

Since its initial release, MGX meanwhile features a wide range of analysis workflows supporting taxonomic as well as functional classification of unassembled metagenome and metatranscriptome datasets. To accompany the novel assembly-based strategies, corresponding workflows have also been implemented for the annotation of assembled contigs and transcripts. These have been carefully tuned to match the characteristics of the read-based analysis workflows in order to facilitate a direct comparison of results.

For functional classification of unassembled metagenome datasets, publicly available analysis resources typically follow a best-hit strategy, *i.e.* the relevant annotation is transferred from the highest-scoring database hit meeting certain predefined quality criteria such as sequence identity, a score, or an E-value cutoff. This approach is valid as the short DNA fragments obtained by sequencing typically do not contain more than one gene fragment or conserved protein domain. For the annotation of assembled contigs and their predicted genes, a different approach is preferable depending on the analysis type. Workflows targeting the annotation of full-length genes will continue to derive the annotation from the best database hit. Pipelines employing protein domain databases such as *e.g.* Pfam are able to identify multiple conserved domains within a single protein-coding sequence. Therefore, these workflows employ a different strategy, where all hits fulfilling the provided quality criteria are retained after an additional step that resolves overlapping predictions.

### MGX user interface components for assembly inspection

Within the MGX user interface, metagenome and metatranscriptome datasets are arranged into groups for joint visualization or statistical evaluation. For MGX 2.0, this central concept has been extended to also support the representation of a dataset partaking in an assembly of one or several datasets. For read-based analyses, the number of sequences assigned to a certain taxon or comprising a gene is simply counted; for annotated assemblies, the computed read coverage of predicted genes is taken into account, as well. Thereby, MGX users are able to define groups for unassembled as well as assembled datasets and directly compare taxonomic classifications or metabolic profiling results in a consistent manner (Fig. 3).

The new “blob plot” component (Fig. 1) enables visual inspection of an assembly, providing a rough overview of an assembled metagenome. The plot is generated interactively and the user is free to select the genomic bins from an assembly to be displayed. Contigs are depicted here as circles in a two-dimensional scatter plot, where the size of each circle correlates with the length of the sequence, and its color is selected based on the taxonomic classification result of the corresponding bin. The individual circles are placed based on the GC content of the assembled contig on the x-axis plotted versus the normalized log-scaled coverage of the sequence on the y-axis.

The “Bin Explorer” module (Fig. 2) facilitates interactive exploration of assembled contigs and their annotations; depending on the currently selected bin, the user can select the contig to be displayed. Contigs are depicted with their predicted genes, and for each gene, a detailed view is available that provides abundance information for this gene in the possibly co-assembled metagenome datasets. Also, annotated subregions are being shown in conjunction with the corresponding annotation data, *e.g.* taxonomic classifications, gene assignments, or detected protein domains.

MGX provides direct access to the DNA or amino-acid sequence of predicted genes, and assembled contigs and transcripts can be exported in standardized formats such as GenBank or FASTA files for downstream processing or deposition in public databases.

### Analysis of public metagenome and metatranscriptome data

We demonstrate the application of the MGX 2.0 software for the analysis of a public microbial community dataset (NCBI SRA project PRJNA321354), which is comprised of both metagenomic and metatranscriptomic sequence data from microbial mats involved in black band disease (BBD) in corals as well as cyanobacterial patches (CP)^39^. Raw sequence data was imported into MGX 2.0 and we performed read-based analyses as well as metagenome and metatranscriptome co-assembly followed by subsequent annotation of predicted genes. Read-based taxonomic profiling was conducted with the MetaPhlAn 4 and Metabuli workflows, as well as with the MGX default classification pipeline^35^. For functional analysis, reads and assembled genes were annotated with EC numbers, Pfam domains, and PGAP HMM models. All workflows were executed applying default parameters suggested by MGX 2.0, which were previously established based on best common practices and internal tool evaluations.

Validation of bin lineages was performed manually with GTDB-Tk (database release 214) applying the *classify wf* workflow (Suppl. Table 5); bin 2 was additionally characterized with the EDGAR platform^40^ (Suppl. Fig. 1).

### Code and Data availability

MGX 2.0 is released for Windows, Linux, and macOS. The full source code is available under the GNU Affero General Public License at https://github. com/MGX-metagenomics; the MGX website with releases and user guide can be found at https://mgx-metagenomics.github.io/. Sequence data used in the evaluation is publicly available from the NCBI SRA under project ID PRJNA321354; the corresponding assemblies and analysis results are offered within the public “MGX2 Demo” project.

## Acknowledgments

Funding for the operation and maintenance of MGX is provided by the German Federal Ministry of Education and Research (BMBF) project “Bielefeld-Gießen Center for Microbial Bioinformatics – BiGi” (grant 031A533) within the German Network for Bioinformatics Infrastructure (de.NBI). Michael R. Crusoe is acknowledged for providing support for the Common Workflow Language (CWL). We gratefully acknowledge all MGX users who submitted bug reports and suggested novel features, and the authors of the black band disease study for making their sequencing data available.

## Contributions

Designed and implemented the software: SJ. Contributed software components: SD. Project funding, outline of development plan and manuscript: AG. Wrote the manuscript: SJ. All authors read and approved the final manuscript.

