## Supplemental material for "MGX 2.0: Shotgun- and assembly-based metagenome and metatranscriptome analysis from a single source"

### Supplementary information

Sebastian Jaenicke*[1,2], Sonja Diedrich[1] & Alexander Goesmann[1]

[1] Bioinformatics and Systems Biology, Justus Liebig University Giessen, Giessen, Germany

[2] ELIXIR Germany, Institute of Bio- and Geosciences (IBG-5) - Computational Metagenomics, Forschungszentrum Jülich GmbH, 52425 Jülich, Germany

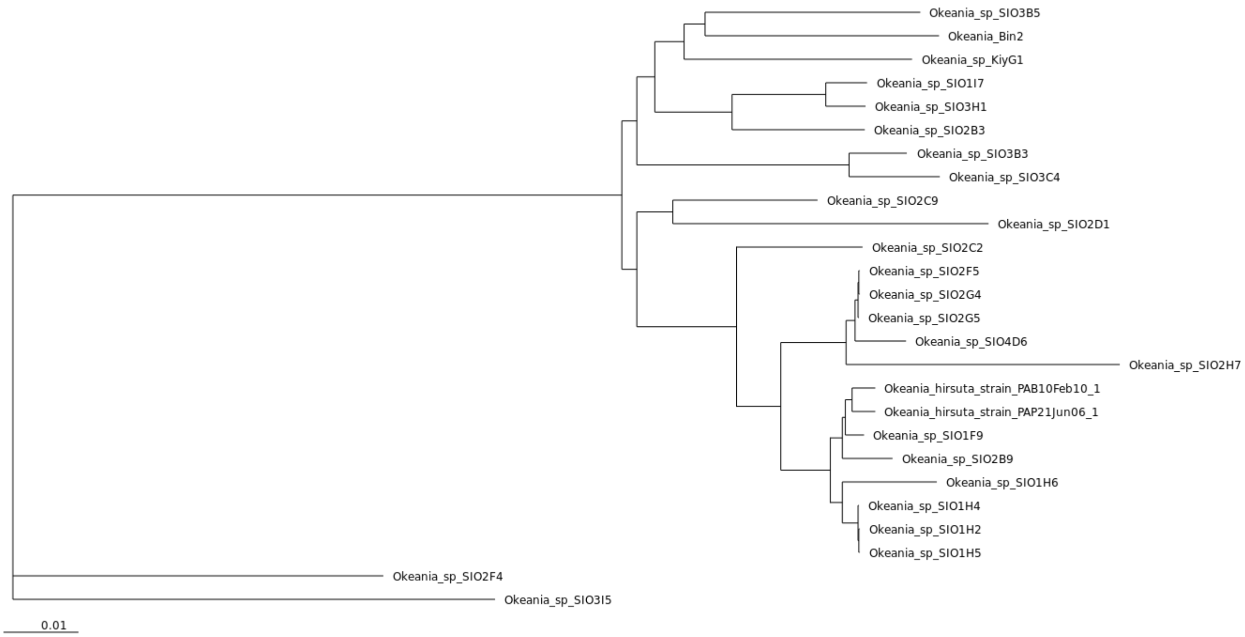


**Suppl. Figure 1: Phylogenetic placement of bin 2.** Assembled sequence data of bin 2 was imported into the EDGAR platform together with publicly available *Okeania* genomes obtained from NCBI. A phylogenetic tree based on the computed core genome confirmed the taxonomic assignment of bin 2 to the *Okeania* genus and close relationship to *Okeania sp. KiyG1*.

**Suppl. Table 1: Workflows for read- and assembly-based taxonomic analysis in MGX 2.0.** With MGX 2.0, a large variety of workflows is provided that allow, in addition to read-based analyses, the annotation of metagenome and metatranscriptome assemblies. All workflows are freely available at https://github.com/MGX-metagenomics/pipelines (Conveyor-based pipelines) and https://github.com/MGX-metagenomics/cwl (CommonWL workflows for metagenome and metatranscriptome assembly).

| Workflow name | read-based | assembly-based |
|---|---|---|
| Centrifuge | ✓ | ✓ |
| Kraken | ✓ | ✗ |
| Kraken 2 | ✓ | ✓ |
| KrakenUniq | ✓ | ✗ |
| Kaiju | ✓ | ✓ |
| Metabuli | ✓ | ✗ |
| MetaPhlAn 4 | ✓ | ✗ |
| MGX default taxonomy | ✓ | ✗ |

**Suppl. Table 2: Workflows for read- and assembly-based functional analysis in MGX 2.0.**

| Workflow name | read-based | assembly-based |
|---|---|---|
| AMRFinderPlus | ✗ | ✓ |
| ARDB | ✓ | ✓ |
| ARG-ANNOT | ✓ | ✗ |
| BacMet | ✓ | ✓ |
| CARD | ✓ | ✓ |
| ClusterMine 360 | ✓ | ✓ |
| dbCAN | ✓ | ✓ |
| EC number | ✓ | ✓ |
| EggNOG | ✓ | ✓ |
| FunGene | ✓ | ✓ |
| KOfam | ✓ | ✓ |
| MIBiG | ✓ | ✓ |
| MVirDB | ✓ | ✓ |
| Pfam | ✓ | ✓ |
| Phage proteins | ✓ | ✓ |
| Plasmid proteins | ✓ | ✗ |
| PGAP | ✓ | ✓ |
| RVDB-prot | ✗ | ✓ |
| TIGRFAM | ✓ | ✓ |
| UniRef50 | ✓ | ✓ |
| VOGDB | ✗ | ✓ |

**Suppl. Table 3: Additional workflows in MGX 2.0.** Additional workflows for reference mapping can be employed to align metagenome and metatranscriptome sequences to reference genomes in order to *e.g.* create fragment recruitment plots or to visually identify gene expression patterns.

| Workflow name | read-based | assembly-based |
|---|---|---|
| Bowtie 2 | ✓ | ✗ |
| FR-HIT | ✓ | ✗ |
| minimap2 | ✓ | ✗ |

**Suppl. Table 4: Improved bin recovery with MGX 2.0.** Taxonomic assignments and CheckM bin assessment for the co-assembly of BBD (SRR3569370) and CP metagenomes (SRR3499156). High quality MAGs (>90% completeness, <5% contamination) in bold.

| Bin | Taxonomy | Contigs | Assembled bp | N50 bp | CDS | Completeness | Contamination |
|---|---|---|---|---|---|---|---|
| Bin 1 | root; Bacteria; Pseudomonadota; Gammaproteobacteria; Oceanospirillales; Oceanospirillaceae | 477 | 2,304,480 | 5,258 | 2,466 | 70.34 | 0.88 |
| **Bin 2** | **root; Bacteria; Cyanobacteriota; Cyanophyceae; Oscillatoriales; Oscillatoriaceae; Okeania; Okeania sp. KiyG1** | 887 | 7,822,907 | 11,248 | 6,120 | 91.16 | 2.4 |
| Bin 3 | root; Bacteria; Pseudomonadota; Alphaproteobacteria; Rhodobacterales; Roseobacteraceae; Shimia | 566 | 2,501,056 | 4,587 | 2,826 | 59.82 | 3.28 |
| Bin 4 | root; Bacteria; Bdellovibrionota; Bdellovibrionia; Bdellovibrionales; Pseudobdellovibrionaceae; Bdellovibrio; Bdellovibrio bacteriovorus | 477 | 1,856,418 | 5,088 | 2,154 | 77.14 | 1.79 |
| Bin 5 | root; Bacteria; Pseudomonadota; Alphaproteobacteria | 819 | 2,666,498 | 4,122 | 3,421 | 71.88 | 2.61 |
| Bin 6 | root; Bacteria; Pseudomonadota; Alphaproteobacteria; Hyphomonadales; Hyphomonadaceae; Hyphomonas; Hyphomonas sp. Mor2 | 570 | 3,198,937 | 6,260 | 3,548 | 77.61 | 2.46 |
| Bin 7 | root; Bacteria; Pseudomonadota; Alphaproteobacteria; Rhodobacterales; Roseobacteraceae; Ruegeria; Ruegeria arenilitoris | 469 | 2,617,389 | 5,952 | 2,896 | 58.63 | 3.33 |
| Bin 8 | root; Bacteria; Bacteroidota; Saprospiria; Saprospirales; Saprospiraceae; Aureispira | 788 | 4,556,000 | 6,558 | 4,304 | 81.91 | 0.25 |
| Bin 9 | root; Bacteria; Chlamydiota; Chlamydiia; Parachlamydiales; Simkaniaceae; Candidatus Neptunochlamydia; Candidatus Neptunochlamydia vexilliferae | 588 | 2,070,345 | 4,559 | 2,261 | 85.88 | 1.42 |
| Bin 10 | root; Bacteria; Pseudomonadota; Gammaproteobacteria | 398 | 3,146,796 | 13,266 | 2,993 | 90.65 | 5.05 |
| Bin 11 | root; Bacteria; Pseudomonadota; Alphaproteobacteria | 419 | 1,808,578 | 4,405 | 2,123 | 49.6 | 1.0 |
| **Bin 12** | **root; Bacteria; Bacteroidota; Cytophagia; Cytophagales** | 170 | 6,512,800 | 66,787 | 5,377 | 99.11 | 1.8 |
| Bin 13 | root; Bacteria; Pseudomonadota; Alphaproteobacteria | 379 | 2,350,285 | 6,964 | 2,554 | 75.93 | 1.17 |
| Bin 14 | root; Bacteria; Pseudomonadota; Alphaproteobacteria; Rhodobacterales; Roseobacteraceae; Ruegeria; Ruegeria arenilitoris | 271 | 3,814,380 | 21,778 | 3,789 | 94.73 | 5.65 |
| Bin 15 | root; Bacteria; Thermodesulfobacteriota; Desulfovibrionia; Desulfovibrionales; Desulfovibrionaceae; Desulfovibrio | 446 | 3,523,557 | 11,523 | 3,439 | 94.61 | 1.45 |
| **Bin 16** | **root; Bacteria; Pseudomonadota; Gammaproteobacteria; Alteromonadales; Alteromonadaceae; Alteromonas** | 90 | 4,244,840 | 72,171 | 3,758 | 96.24 | 0.37 |
| **Bin 17** | **root; Bacteria; Bacteroidota; Cytophagia; Cytophagales; Microscillaceae; Microscilla; Microscilla marina** | 269 | 6,276,528 | 36,430 | 6,101 | 95.91 | 2.17 |
| **Bin 18** | **root; Bacteria; Bacteroidota; Cytophagia; Cytophagales; Reichenbachiellaceae; Ekhidna; Ekhidna lutea** | 165 | 3,622,682 | 36,025 | 3,323 | 98.21 | 0.71 |
| Bin 19 | root; Bacteria; Bdellovibrionota; Bdellovibrionia; Bdellovibrionales; Pseudobdellovibrionaceae; Micavibrio; Micavibrio aeruginosavorus | 114 | 1,931,895 | 26,687 | 1,990 | 92.96 | 2.45 |
| **Bin 20** | **root; Bacteria; Bacteroidota; Flavobacteriia; Flavobacteriales; Flavobacteriaceae; Winogradskyella; Winogradskyella psychrotolerans** | 108 | 3,160,775 | 64,586 | 2,899 | 98.35 | 1.49 |
| **Bin 21** | **root; Bacteria; Bacteroidota; Flavobacteriia; Flavobacteriales; Flavobacteriaceae; Winogradskyella** | 115 | 3,534,206 | 48,721 | 3,076 | 98.76 | 0.29 |
| **Bin 22** | **root; Bacteria; Pseudomonadota; Alphaproteobacteria; Rhodobacterales; Roseobacteraceae; Ruegeria; Ruegeria arenilitoris** | 46 | 3,277,785 | 107,437 | 3,202 | 100.0 | 0.43 |
| **Bin 23** | **root; Bacteria; Campylobacterota; Epsilonproteobacteria; Campylobacterales; Arcobacteraceae; Arcobacter; Arcobacter sp. LA11** | 107 | 3,018,567 | 49,339 | 3,028 | 99.59 | 0.27 |
| **Bin 24** | **root; Bacteria; Cyanobacteriota; Cyanophyceae; Desertifilales; Desertifilaceae; Roseofilum; Roseofilum reptotaenium** | 108 | 5,486,743 | 79,582 | 4,869 | 99.11 | 0.22 |
| **Bin 25** | **root; Bacteria; Pseudomonadota; Gammaproteobacteria; Oceanospirillales; Oceanospirillaceae; Bacterioplanoides; Bacterioplanoides sp. SCSIO 12839** | 37 | 3,885,291 | 213,889 | 3,575 | 99.57 | 1.02 |
| Unbinned | root; | 45,718 | 95,586,388 | 2,128 | 110,708 | 100.0 | 1839.11 |

**Suppl. Table 5: GTDB-Tk lineages for assembled bins.** GTDB taxonomy assignments were computed with `gtdbtk classify_wf` outside of MGX 2.0 using database version 214.

| Bin | GTDB assignment |
|---|---|
| Bin 1 | d__Bacteria;p__Pseudomonadota;c__Gammaproteobacteria;o__Pseudomonadales;f__DSM-6294;g__Oceanobacter;s__ |
| Bin 2 | d__Bacteria;p__Cyanobacteriota;c__Cyanobacteriia;o__Cyanobacteriales;f__Microcoleaceae;g__Okeania;s__ |
| Bin 3 | d__Bacteria;p__Pseudomonadota;c__Alphaproteobacteria;o__Rhodobacterales;f__Rhodobacteraceae;g__Shimia;s__ |
| Bin 4 | d__Bacteria;p__Bdellovibrionota;c__Bdellovibrionia;o__Bdellovibrionales;f__JAMLIO01;g__JALZUR01;s__ |
| Bin 5 | d__Bacteria;p__Pseudomonadota;c__Alphaproteobacteria;o__UBA2562;f__UBA2562;g__;s__ |
| Bin 6 | d__Bacteria;p__Pseudomonadota;c__Alphaproteobacteria;o__Caulobacterales;f__Hyphomonadaceae;g__Henriciella;s__ |
| Bin 7 | d__Bacteria;p__Pseudomonadota;c__Alphaproteobacteria;o__Rhodobacterales;f__Rhodobacteraceae;g__Ruegeria;s__ |
| Bin 8 | d__Bacteria;p__Bacteroidota;c__Bacteroidia;o__Chitinophagales;f__Saprospiraceae;g__Aureispira;s__ |
| Bin 9 | d__Bacteria;p__Chlamydiota;c__Chlamydiia;o__Chlamydiales;f__Simkaniaceae;g__Neptunochlamydia;s__ |
| Bin 10 | d__Bacteria;p__Pseudomonadota;c__Gammaproteobacteria;o__Enterobacterales_A;f__Neiellaceae;g__;s__ |
| Bin 11 | d__Bacteria;p__Pseudomonadota;c__Alphaproteobacteria;o__UBA2562;f__UBA2562;g__;s__ |
| Bin 12 | d__Bacteria;p__Bacteroidota;c__Bacteroidia;o__Cytophagales;f__Cyclobacteriaceae;g__JANSWY01;s__ |
| Bin 13 | d__Bacteria;p__Pseudomonadota;c__Alphaproteobacteria;o__UBA2562;f__UBA2562;g__;s__ |
| Bin 14 | d__Bacteria;p__Pseudomonadota;c__Alphaproteobacteria;o__Rhodobacterales;f__Rhodobacteraceae;g__JABSSB01;s__ |
| Bin 15 | d__Bacteria;p__Desulfobacterota;c__Desulfovibrionia;o__Desulfovibrionales;f__Desulfovibrionaceae;g__Maridesulfovibrio;s__ |
| Bin 16 | d__Bacteria;p__Pseudomonadota;c__Gammaproteobacteria;o__Enterobacterales_A;f__Alteromonadaceae;g__Alteromonas;s__ |
| Bin 17 | d__Bacteria;p__Bacteroidota;c__Bacteroidia;o__Cytophagales;f__Microscillaceae;g__;s__ |
| Bin 18 | d__Bacteria;p__Bacteroidota;c__Bacteroidia;o__Cytophagales;f__Cyclobacteriaceae;g__Ekhidna;s__ |
| Bin 19 | d__Bacteria;p__Pseudomonadota;c__Alphaproteobacteria;o__Micavibrionales;f__Micavibrionaceae;g__TMED27;s__ |
| Bin 20 | d__Bacteria;p__Bacteroidota;c__Bacteroidia;o__Flavobacteriales;f__Flavobacteriaceae;g__DT-34;s__ |
| Bin 21 | d__Bacteria;p__Bacteroidota;c__Bacteroidia;o__Flavobacteriales;f__Flavobacteriaceae;g__Winogradskyella;s__ |
| Bin 22 | d__Bacteria;p__Pseudomonadota;c__Alphaproteobacteria;o__Minwuiales;f__;g__;s__ |
| Bin 23 | d__Bacteria;p__Campylobacterota;c__Campylobacteria;o__Campylobacterales;f__Arcobacteraceae;g__Halarcobacter;s__ |
| Bin 24 | d__Bacteria;p__Cyanobacteriota;c__Cyanobacteriia;o__Cyanobacteriales;f__Desertifilaceae;g__Roseofilum;s__Roseofilum |
| Bin 25 | d__Bacteria;p__Pseudomonadota;c__Gammaproteobacteria;o__Pseudomonadales;f__DSM-6294;g__Bacterioplanoides;s__ |

**Suppl. Table 6: Software versions used within the metagenome and metatranscriptome assembly workflows.** Compile options and additional patches are available at https://github.com/MGX-metagenomics/tools.

| Tool | Version |
|---|---|
| assignBin | 1.0 |
| bamstats | 1.0 |
| CheckM | 1.2.2 |
| DAS Tool | 1.1.6 |
| fastp | 0.23.2 |
| featureCounts | 2.0.6 |
| MetaBAT2 | 2.16-4-g40efa2d |
| Metabuli | 1.0.1 |
| MEGAHIT | 1.2.9 |
| prodigal | 2.6.3 |
| RiboDetector | 0.2.7 |
| rnaSPAdes | 3.15.5 |
| samtools | 1.18 |
| SemiBin2 | 1.5.1 |
| strobealign | 0.11.0 |
| Trimmomatic | 0.39 |
| Vamb | 4.1.3 |

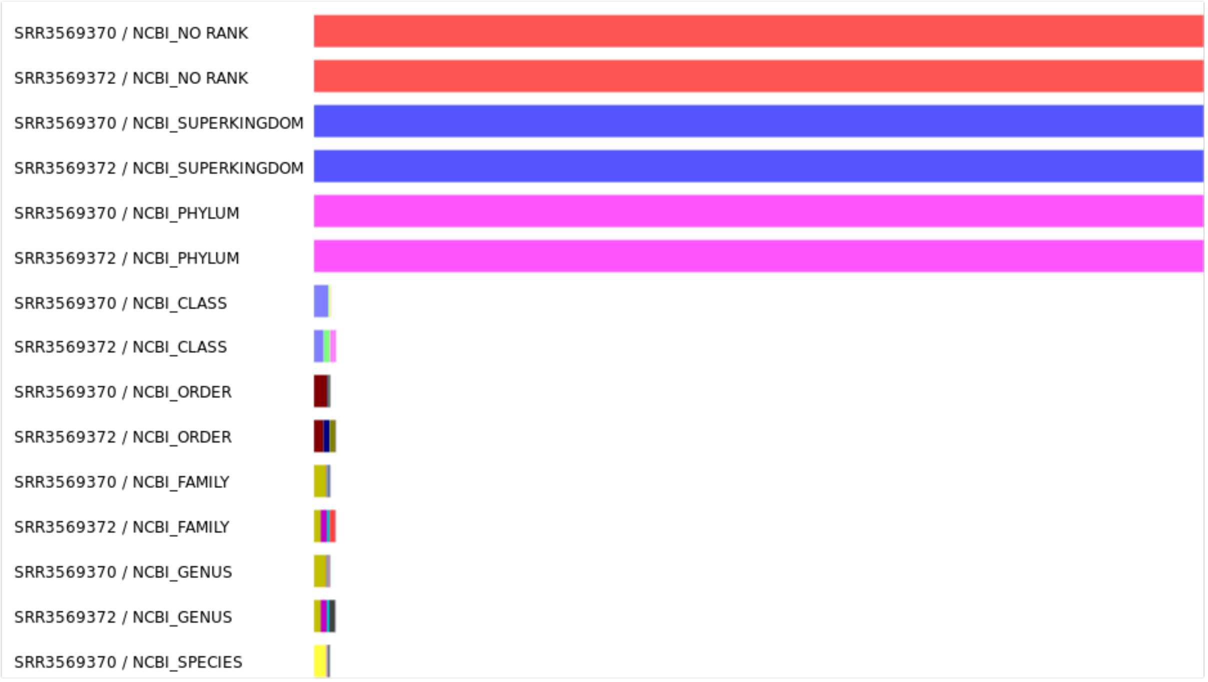


**Suppl. Figure 2: MetaPhlAn 4 taxonomic profiling results of the BBD metagenome and transcriptome.** Unassembled reads of the BBD metagenome (SRR3569370) and metatranscriptome (SRR3569372) were taxonomically profiled with the MetaPhlAn workflow within MGX (MetaPhlAn 4.0.6; database version Oct22). Data normalized to root rank. Apparent is a largely reduced number of taxonomic assignments below the phylum rank.

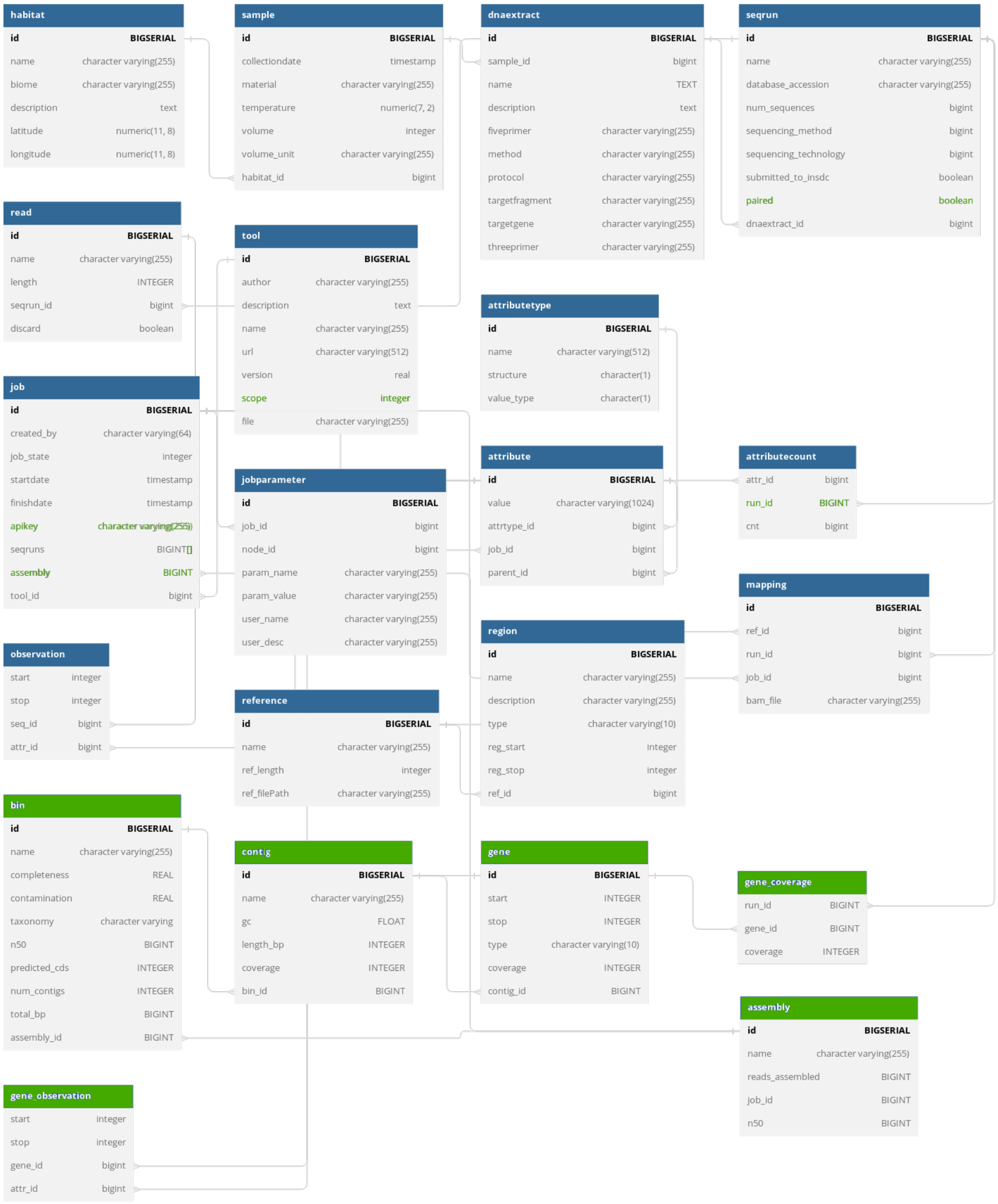


**Suppl. Figure 3: MGX 2.0 database schema.** The PostgreSQL database schema was extended with additional tables to store information about an assembly including bins, contigs, and predicted genes. For each gene, read coverage is computed separately for each dataset partaking in an assembly, thereby allowing to recover abundance information. New database entities and changed attributes are shown in green. Diagram created with https://dbdiagram.io.
